# Jumping To Conclusions, General Intelligence, And Psychosis Liability: Findings From The Multi-Centre EU-GEI Case-Control Study

**DOI:** 10.1101/634352

**Authors:** Giada Tripoli, Diego Quattrone, Laura Ferraro, Charlotte Gayer-Anderson, Victoria Rodriguez, Caterina La Cascia, Daniele La Barbera, Crocettarachele Sartorio, Ilaria Tarricone, Domenico Berardi, Andrei Szöke, Celso Arango, Andrea Tortelli, Pierre-Michel Llorca, Lieuwe de Haan, Eva Velthorst, Julio Bobes, Miguel Bernardo, Julio Sanjuán, Jose Luis Santos, Manuel Arrojo, Cristina Marta Del-Ben, Paulo Rossi Menezes, Jean-Paul Selten, EU-GEI WP2 Group, Peter B Jones, Alex Richards, Michael O’Donovan, Bart PF Rutten, Jim van Os, Craig Morgan, Pak C Sham, Robin M Murray, Graham K Murray, Marta Di Forti

## Abstract

**Background:** The “jumping to conclusions” (JTC) bias is associated with both psychosis and general cognition but their relationship is unclear. In this study, we set out to clarify the relationship between the JTC bias, IQ, psychosis and polygenic liability to schizophrenia and IQ.

**Methods:** 817 FEP patients and 1294 population-based controls completed assessments of general intelligence (IQ), and JTC (assessed by the number of beads drawn on the probabilistic reasoning “beads” task) and provided blood or saliva samples from which we extracted DNA and computed polygenic risk scores for IQ and schizophrenia.

**Results:** The estimated proportion of the total effect of case/control differences on JTC mediated by IQ was 79%. Schizophrenia Polygenic Risk Score (SZ PRS) was non-significantly associated with a higher number of beads drawn (B= 0.47, 95% CI −0.21 to 1.16, p=0.17); whereas IQ PRS (B=0.51, 95% CI 0.25 to 0.76, p<0.001) significantly predicted the number of beads drawn, and was thus associated with reduced JTC bias. The JTC was more strongly associated with higher level of psychotic-like experiences (PLE) in controls, including after controlling for IQ (B= −1.7, 95% CI −2.8 to −0.5, *p*=0.006), but did not relate to delusions in patients.

**Conclusions:** the JTC reasoning bias in psychosis is not a specific cognitive deficit but is rather a manifestation or consequence, of general cognitive impairment. Whereas, in the general population, the JTC bias is related to psychotic-like experiences, independent of IQ. The work has potential to inform interventions targeting cognitive biases in early psychosis.

## Introduction

Jumping to conclusions (JTC) is a well-established reasoning and data gathering bias found in patients with psychosis (Dudley et al, 2016; Garety & Freeman, 2013; So et al, 2016). It is usually measured by a reasoning task based on a Bayesian model of probabilistic inference known as the “beads task” (Dudley et al, 1997; Huq et al, 1988).

Early work investigating reasoning bias through beads in jars paradigm focused on the association with delusions, finding that patients with delusions tend to require fewer beads to reach a decision than would be expected following Bayesian norms (Huq et al, 1988). In this respect, it has been suggested that data-gathering bias represents a key cognitive component in delusion formation and maintenance (Garety & Freeman, 1999). Although most of the studies endorsing this association used a cross-sectional design, comparing those with and without delusions in samples of patients with schizophrenia or other psychoses (Freeman et al, 2014; Garety et al, 2013), meta-analyses mostly found a weak association with delusions, yet a clear association with psychosis (Dudley et al, 2016; Fine et al, 2007; Ross et al, 2015; So et al, 2016). Moreover, jumping to conclusions was found not only in non-delusional and remitted patients with schizophrenia, but also in individuals at clinical high risk and first-degree relatives (Broome et al, 2007; Menon et al, 2006; Moritz & Woodward, 2005; Peters & Garety, 2006; Van Dael et al, 2005).

Overall, these findings suggest that JTC could play a role in the liability for psychotic disorders, perhaps serving as an endophenotype associated with the genetic risk. As a highly heritable and polygenic disorder, many common genetic variants contribute to the risk of schizophrenia, which can be summarised into an individual polygenic risk score (PRS) (International Schizophrenia Consortium, 2009; Schizophrenia Working Group of the Psychiatric Genomics, 2014). GWAS studies have found a significant genetic overlap between cognition and schizophrenia (Mistry et al, 2018; Ohi et al, 2018; Shafee et al, 2018), and a recent study by Toulopoulou (Toulopoulou et al, 2018) showed that the variance in schizophrenia liability explained by PRS was partially mediated through cognitive deficit. Despite a large body of literature on the JTC bias describing it as a potential intermediate phenotype, to date there has been no investigation into the possible genetic overlap with psychosis through PRS strategy. Nonetheless, there is a considerable amount of literature suggesting a link between JTC and cognitive functions. Cross-sectional studies on both recent onset psychosis and schizophrenia suggest that patients who present with neuropsychological deficits, especially involving executive functions, display more tendency to jump to conclusions (Garety et al, 2013; González et al, 2018). Those findings appear to be corroborated by the study of Lunt (Lunt et al, 2012) where JTC bias was found to be more prominent in individuals with prefrontal lesions, especially in the left side of cortex, compared with controls. Woodward (Woodward et al, 2009), using a principal component analysis in a sample of inpatients with schizophrenia found that neuropsychological functions and JTC loaded on the same factor. Similar findings were obtained in a confirmatory factor analysis on psychotic patients with delusions and controls (Bentall et al, 2009). Interestingly in the latter, when general cognitive functioning was included in the model, the association between paranoia and JTC no longer held. In their study comparing patients with delusions, remitted patients with schizophrenia, and controls, Lincoln (Lincoln et al, 2010) found that, after taking intelligence into account, not only the association between JTC and delusions disappeared, but the group effect on the JTC bias was not detected any longer. Likewise, general intelligence affected statistically the relation between JTC and psychosis liability in the van Dael study (Van Dael et al, 2005). Lower IQ was also found associated with JTC in First Episode Psychosis (Catalan et al, 2015; Falcone et al, 2015). Nonetheless, other studies on early psychosis did not detect any association with cognition, possibly due to small sample size (Langdon et al, 2014; Ormrod et al, 2012).

Thus, although many studies highlighted that jumping to conclusions is strongly associated with delusions, and more broadly with psychosis, this association seems to be affected by general cognitive function. To address this question robustly requires a large sample size, whereas most previous studies of JTC in psychiatry have had moderate sample sizes of between 20 and 100 patients. Therefore, we used data from the large multi-country EUropean Network of national schizophrenia networks studying Gene-Environment Interactions (EUGEI) case-control study of first episode psychosis to investigate whether IQ play a role as mediator in the pathway between the bias and the disorder. We also aimed to investigate whether JTC is associated with the liability for psychotic disorders and/or liability to general intellectual function.

Therefore, in this study we aim to test the following predictions:

1. The JTC bias will be best predicted by clinical status through IQ mediation;
2. The JTC bias will be predicted by the Polygenic Risk Score for Schizophrenia (SZ PRS) and by the Polygenic Risk Score for IQ (IQ PRS).
3. The JTC bias will be predicted by delusions in patients and psychotic-like experiences (PLE) in controls.

## Methods

### Participants

Participants were recruited and assessed as part of the incidence and first episode case-control study, conducted as part of the EU-GEI programme (Gayer-Anderson et al, *in submission*; Jongsma et al, 2018). The study was designed to investigate risk factors for psychotic disorders over three-year period between 1/5/2010 and 1/4/2015 in 17 catchment areas in England, France, the Netherlands, Italy, Spain and Brazil.

All participants provided informed, written consent following full explanation of the study. Ethical approval was provided by relevant research ethics committees in each of the study sites. Patients with a first episode psychosis (FEP) were recruited through regular checks across the 17 defined catchment areas Mental Health services to identify all individuals aged 18 – 64 years who presented with a first episode psychosis during the study period. Patients were included if they met the following criteria during the recruitment period: (a) aged between 18 and 64 years; (b) presentation with a clinical diagnosis for an untreated FEP, even if longstanding (International Statistical Classification of Diseases and Related Health Problems, Tenth Revision [ICD-10] codes F20-F33); (c) resident within the catchment area at FEP. Exclusion criteria were: (a) previous contact with psychiatric services for psychosis; (b) psychotic symptoms with any evidence of organic causation; and (c) transient psychotic symptoms resulting from acute intoxication (ICD-10: F1*x*.5).

Inclusion criteria for controls were: a) aged between 18 and 64 years; b) resident within a clearly defined catchment area at the at the time of consent into the study; c) sufficient command of the primary language at each site to complete assessments; and d) no current or past psychotic disorder. To select a population-based sample of controls broadly representative of local populations in relation to age, gender, and ethnicity, a mixture of random and quota sampling was adopted. Quotas for control recruitment were based on the most accurate local demographic data available, and then filled using a variety of recruitment methods, including through: 1) random sampling from lists of all postal addresses (e.g., in London); 2) stratified random sampling via GP lists (e.g., in London and Cambridge) from randomly selected surgeries; and 3) ad hoc approaches (e.g., internet and newspaper adverts, leaflets at local stations, shops, and job centres).

### Measures

Detailed information on age, sex, self-reported ethnicity, level of education and social networks was collected using the Medical Research Council (MRC) Sociodemographic Schedule (Mallett, 1997).

The short form of the WAIS III was administered as an indicator of general cognitive ability (IQ) and includes the following subtests: Information (verbal comprehension), Block Design (reasoning and problem solving), Arithmetic (working memory) and Digit symbol-coding (processing speed). This short form has been shown to give reliable and valid estimates of Full-Scale IQ in schizophrenia (Velthorst et al, 2013).

Following instructions and practice, the JTC bias was assessed using a single trial of the computerised 60:40 version of the beads task. In this task, there are two jars containing coloured beads with complementary ratio (60:40). One jar is chosen, and beads are drawn one at time and shown to the participants following the pattern: BRRBBRBBBRBBBBRRBRRB, where B indicates a blue bead and R indicates red. After each draw they are required to either decide from which jar the beads have come or defer the decision (up to a maximum of 20 beads). This single-trial experiment was terminated when the participant made the decision of which jar the beads were being drawn from. The key outcome variable employed as an index of the JTC bias was the number of “Draws-To-Decision” (DTD); the lower the DTD, the greater the JTC bias.

Psychopathology was assessed using the OPerational CRITera system (McGuffin et al, 1991). Item response modelling was previously used to develop a bi-factor model composed of general and specific dimensions of psychotic symptoms, which include the positive symptom dimension (delusion and hallucination items) (Quattrone et al, 2018). For the purposes of the present work, we adapted the previous method to estimate an alternative bi-factor model comprising two discrete hallucination and delusion symptom dimensions, instead of a single positive symptom dimension. We assessed psychotic-like experiences (PLE) in controls through the Community Assessment of Psychic Experience (CAPE) (Stefanis et al, 2017) (http://www.cape42.homestead.com/) positive dimension score (Supplementary Table S1).

### Polygenic Risk Scores

The case-control genotyped WP2 EUGEI sample (N=2,169; cases’ samples N=920, controls’ samples N=1,248) included DNA extracted from blood (N=1,857) or saliva (N=312). The samples were genotyped at the MRC Centre for Neuropsychiatric Genetics and Genomics in Cardiff (UK) using a custom Illumina HumanCoreExome-24 BeadChip genotyping array covering 570,038 genetic variants. After genotype Quality Control, we excluded SNPs with minor allele frequency <0.5%, Hardy Weinberg Equilibrium p<10-6, missingness >2%. After sample Quality Control, we excluded samples with >2% missingness, heterozygosity Fhet >0.14 or <-0.11, who presented genotype-phenotype sex mismatch or who clustered with homogenous black ancestry in PCA analysis (N=170). The final sample was composed of 1,720 individuals (1,112 of European ethnicity, 608 of any other ethnicities but not black African), of which 1,041 controls and 679 patients. Imputation was performed through the Michigan Imputation Server, using the Haplotype Reference Consortium reference panel with the Eagle software for estimating haplotype phase, and Minimac3 for genotype imputation (Das et al, 2016; Loh et al, 2016; McCarthy et al, 2016). The imputed variants with r2 < 0.6, MAF < 0.1% or missingness > 1% were excluded.

The polygenic risk score for schizophrenia and IQ were built using, as training data sets, the results from the last available mega-analysis from the Psychiatric Genomics Consortium (PGC)(Schizophrenia Working Group of the Psychiatric Genomics, 2014) and Savage et al(Savage et al, 2018) respectively. In PRSice, individuals’ number of risk alleles in the target sample were weighted by the log odds ratio from the discovery samples and summed into the PRSs at a 0.05 SNPs Pt-thresholds (apriori selected). We excluded people of homogeneous African ancestry since in this population the SZ PRS from the PGC2 we calculated, as reported by other studies(Vassos et al, 2017), failed to explain a significant proportion of the variance (R^2^=1.1%, *p*=0.004).

### Design and Procedure

The EU-GEI study WP2 employed a case-control design collecting data with an extensive battery of demographic, clinical, social, and biological measures (Core assessment); psychological measures, and cognitive tasks. All EU-GEI WP2 participants with JTC and IQ data were included in the current study. All the researchers involved in administrating the assessments undertook a training organised by a technical working committee of the overall EU-GEI study (Work Package 11) at the beginning and throughout the study. Inter-rater reliability was assessed annually to warrant comparability of procedures and methods across sites.

### Statistical analysis

Analyses were conducted in STATA 15 (StataCorp., 2017). Preliminary descriptive analyses were performed using chi-square and t tests to examine the differences on age, sex, ethnicity, level of education, IQ, and DTD between cases and controls. To test possible statistical mediation by IQ as intervening variable between case/control status and DTD, we applied Baron and Kenny’s procedure (1986). According to the authors, a mediation can be established when 1) variations in the independent variable (IV) significantly account for variations in the dependent variable (DV); 2) variations in the IV significantly account for variations in the mediator variable (MV); 3) variations in the MV significantly account for variations in the DV; 4) a previous significant effect of IV on DV is markedly decreased after controlling for MV. Perfect mediation occurs when the independent variable has no effect after controlling for the mediator (Baron & Kenny, 1986).

The STATA 15 *sgmediation* command was used to perform three OLS regressions according to Baron and Kenny’s steps as follows: 1) DTD (DV) regressed on case/control status (IV) – Path c, 2) IQ (MV) regressed on case/control status (IV) – Path a, 3) DTD (DV) regressed on IQ (MV) and case/control status (IV) – Path b and c’ (Figure 1). All the steps included as covariates age, sex, ethnicity, and country. Furthermore, to generate confidence intervals for the indirect effect, 5000 bootstrap replications were performed (Preacher & Hayes, 2008) and non-parametric confidence interval based on the empirical sampling distributions was constructed.

**Figure 1.**
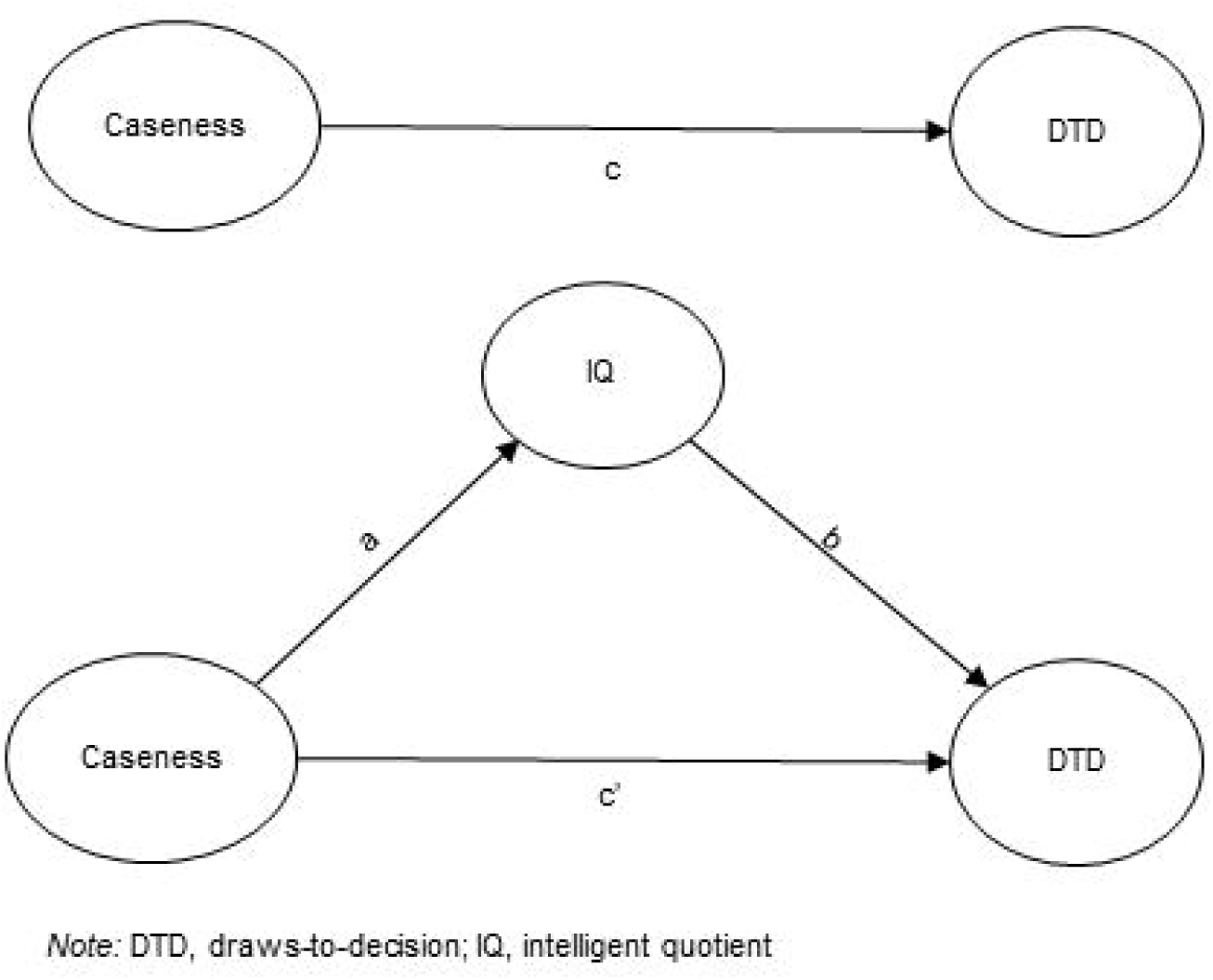
Mediation model between Caseness (IV), IQ(MV), and DTD(DV)

To investigate whether jumping to conclusions was associated with the liability for psychotic disorders and general intelligence, we firstly tested the accuracy of the PRSs to predict their primary phenotypes (case/control status and IQ) in our sample through logistic and linear regression models respectively. Then, we built and compared linear regression models regressing DTD on 1) case/control status, controlling for age, sex, and 20 principal components for population stratification; 2) SZ PRS and 3) IQ PRS, adjusting for case/control status, age, sex, and 20 principal components for population stratification.

Linear regression models were built to test the effect of the delusion symptom dimension on DTD, adding in a second step as covariates age, sex, ethnicity, IQ, and country, in patients. We ran the same models in controls using the CAPE positive score as a predictor.

## Results

### Sample characteristics

Cases and controls recruited as part of the EU-GEI study were included in the current study if data on both Beads task and WAIS were available. This led to a sample of FEP cases=817 and controls N=1,294 for the mediation analysis (Figure 2).

**Figure 2.**
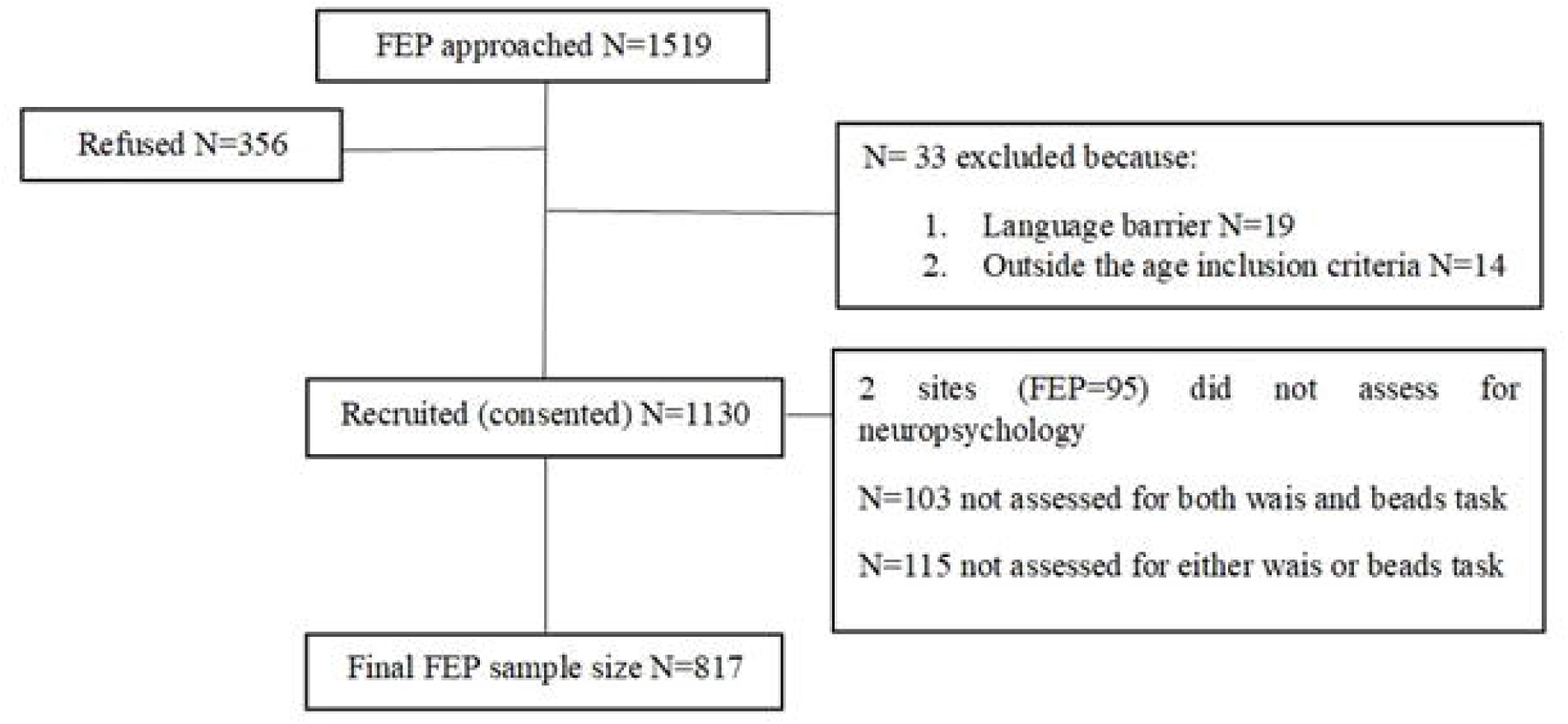
FEP recuitment flowchart.

Cases were more often younger (cases mean age=30.6±10.4 vs. controls mean age=36.2±13.1; t=10.3, p<0.001), men [cases 61.2% (500) vs. controls 47.5 (615); χ^2^_(1)_=37. 6, p<0.001] and from minority ethnic backgrounds (χ^2^_(6)_= 48.3, p<0.001) compared to controls. Controls were more likely to have achieved higher level of education (χ^2^_(3)_=202.5, p<0.001) than cases (Table 1). The aforementioned differences are those expected when comparing psychotic patients with the general population.

**Table 1.**
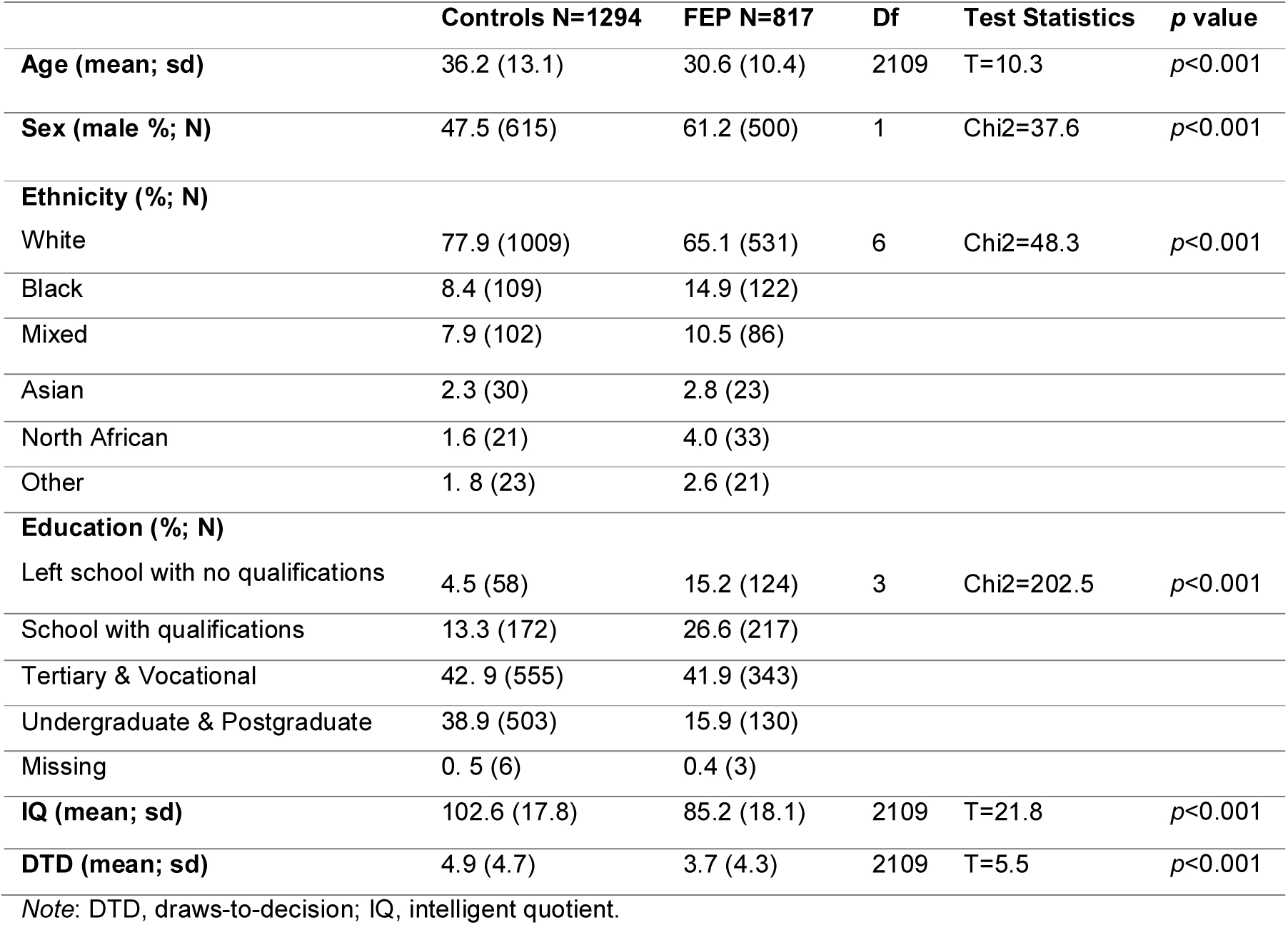
Demographic and cognitive characteristics of the sample included in the analysis.

|  | **Controls N=1294** | **FEP N=817** | **Df** | **Test Statistics** | **p value** |
| --- | --- | --- | --- | --- | --- |
| **Age (mean; sd)** | 36.2 (13.1) | 30.6 (10.4) | 2109 | T=10.3 | $p<0.001$ |
| **Sex (male %; N)** | 47.5 (615) | 61.2 (500) | 1 | Chi2=37.6 | $p<0.001$ |
| **Ethnicity (%; N)** |  |  |  |  |  |
| White | 77.9 (1009) | 65.1 (531) | 6 | Chi2=48.3 | $p<0.001$ |
| Black | 8.4 (109) | 14.9 (122) |  |  |  |
| Mixed | 7.9 (102) | 10.5 (86) |  |  |  |
| Asian | 2.3 (30) | 2.8 (23) |  |  |  |
| North African | 1.6 (21) | 4.0 (33) |  |  |  |
| Other | 1.8 (23) | 2.6 (21) |  |  |  |
| **Education (%; N)** |  |  |  |  |  |
| Left school with no qualifications | 4.5 (58) | 15.2 (124) | 3 | Chi2=202.5 | $p<0.001$ |
| School with qualifications | 13.3 (172) | 26.6 (217) |  |  |  |
| Tertiary & Vocational | 42.9 (555) | 41.9 (343) |  |  |  |
| Undergraduate & Postgraduate | 38.9 (503) | 15.9 (130) |  |  |  |
| Missing | 0.5 (6) | 0.4 (3) |  |  |  |
| **IQ (mean; sd)** | 102.6 (17.8) | 85.2 (18.1) | 2109 | T=21.8 | $p<0.001$ |
| **DTD (mean; sd)** | 4.9 (4.7) | 3.7 (4.3) | 2109 | T=5.5 | $p<0.001$ |

*Note:* DTD, draws-to-decision; IQ, intelligent quotient.

Table 1 also shows both IQ (cases mean=85.2±18.1 vs. controls mean=102.6±17.8, t=21.8, p<0.001) and DTD (cases mean=3.7±4.3 vs. controls mean=4.9±4.7, t=5.5, p<0.001) were lower in cases compared to controls.

The analysis on PRSs was performed in a subsample of 519 FEP and 881 population controls with available GWAS data.

### Jumping to Conclusions, psychosis and IQ

The results of the mediation model are displayed in Figure 3. There was evidence of a negative association between caseness and draws to decision in path c [B= −1.4 (0.2); 95% CI: −1.8 to −0.9; *p*<0.001]; as well as with IQ in Path a [B= −16. 7 (0.8); 95% CI: −18.24 to - 15.10; *p*<0.001]. Path b showed that higher IQ scores were significantly related to more draws to decision [B=0.06 (0.01); 95% CI: 0.05 to 0.07; *p*<0.001]. The direct effect of case status on DTD displayed in path c’ markedly dropped from −1.4 to −0.3 and was no longer statistically significant [B= −0.3 (0.2); 95% CI: −0.7 to 0.1; *p*=0.19]. The estimated proportion of the total effect of case status on DTD mediated by IQ was 79%. After performing 5000 bootstrap replications, the estimated indirect effect remained statistically significant [B= −0.2 (0.02); 95% CI: −0.3 to −0.2].

**Figure 3.**
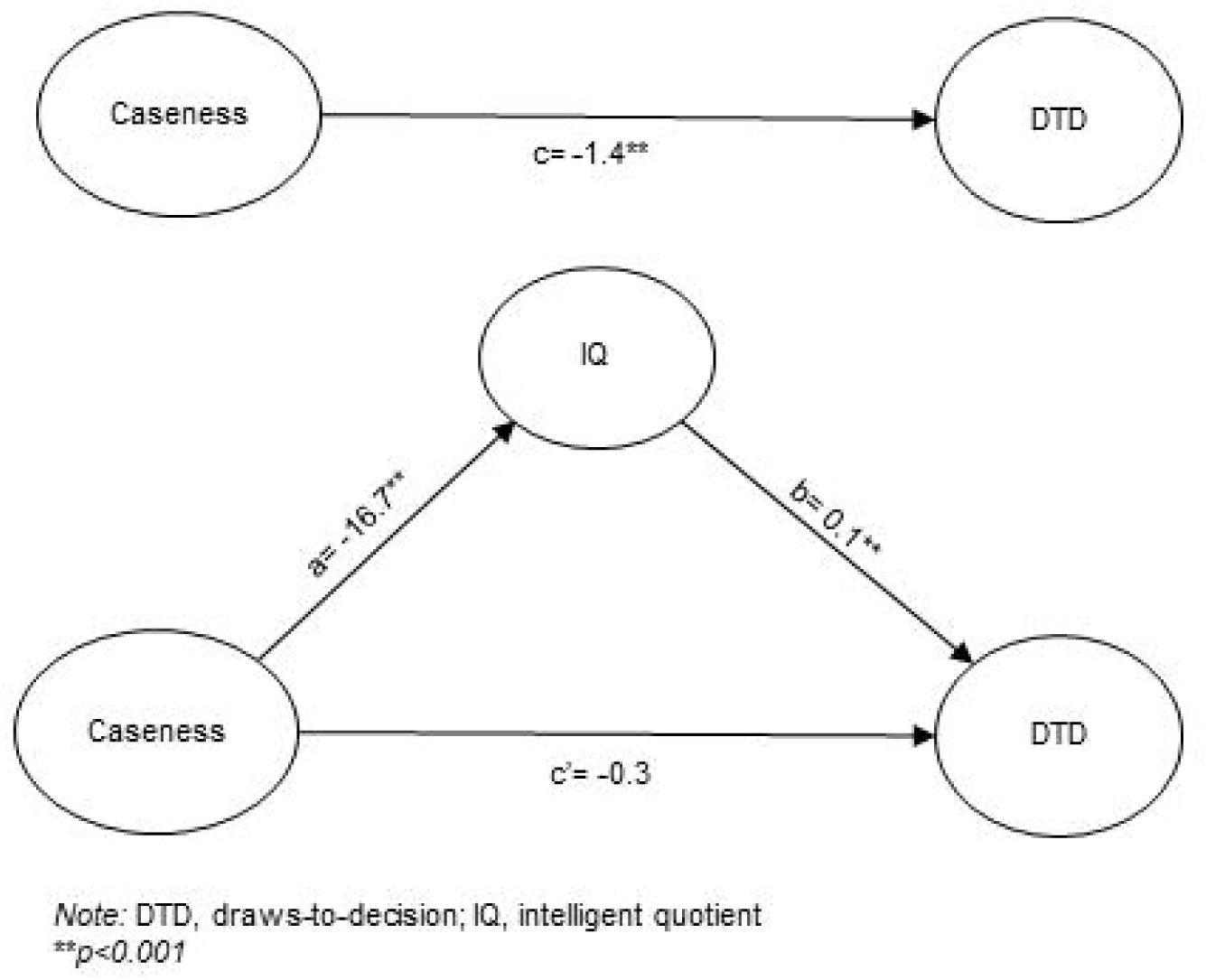
Mediation results.

### Jumping to Conclusions, SZ PRS, and IQ PRS

SZ PRS was a predictor for case status (OR=5.3, 95% CI 3.7 to 7.5, *p*<0.001), explaining 7% of the variance, as well as IQ PRS for IQ (B=3.9, 95% CI 2.9 to 4.9, *p*<0.001), accounting for 14% of the variance. As it is shown in table 2, SZ PRS was not significantly associated with jumping to conclusions, but in fact was non-significantly association with increased numbers of beads drawn (B=0.5, 95% CI −0.2 to 1.2, *p*=0.17); whereas IQ PRS was positively associated with the number of beads drawn (B=0.5, 95% CI 0.3 to 0.8, *p*<0.001). When adding the IQ PRS to case-control status as a predictor of DTD, the variance explained rose to 10%, of which 1% was due to the PRS term. Since the SZ PRS failed to predict DTD, violating the first assumption for a mediation, the further mediation analysis was not pursued. The relationship between DTD and delusions is reported in the supplement.

**Table 2.**
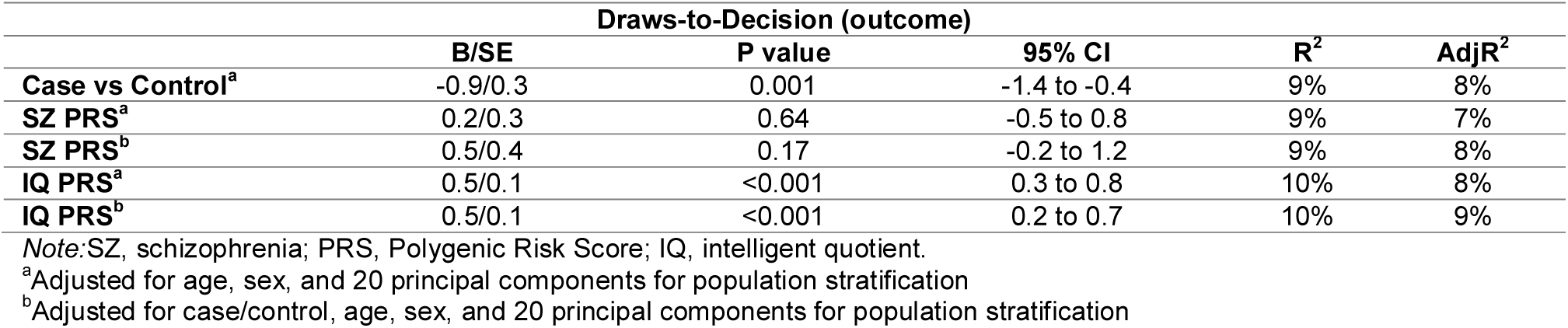
Linear Regressions of Polygenic Risk Scores Predicting DTD.

|  | Draws-to-Decision (outcome) |  |  |  |  |
| --- | --- | --- | --- | --- | --- |
|  | B/SE | P value | 95% CI | R<sup>2</sup> | AdjR<sup>2</sup> |
| Case vs Control<sup>a</sup> | -0.9/0.3 | 0.001 | -1.4 to -0.4 | 9% | 8% |
| SZ PRS<sup>a</sup> | 0.2/0.3 | 0.64 | -0.5 to 0.8 | 9% | 7% |
| SZ PRS<sup>b</sup> | 0.5/0.4 | 0.17 | -0.2 to 1.2 | 9% | 8% |
| IQ PRS<sup>a</sup> | 0.5/0.1 | <0.001 | 0.3 to 0.8 | 10% | 8% |
| IQ PRS<sup>b</sup> | 0.5/0.1 | <0.001 | 0.2 to 0.7 | 10% | 9% |

Note: SZ, schizophrenia; PRS, Polygenic Risk Score; IQ, intelligent quotient.
<sup>a</sup>Adjusted for age, sex, and 20 principal components for population stratification
<sup>b</sup>Adjusted for case/control, age, sex, and 20 principal components for population stratification

### Jumping to conclusions, positive symptoms, and psychotic-like experiences

The delusion symptom dimension was *positively* associated with draws to decision (B=0.4, 95% CI 0.1 to 0.8, p=0.013), although the association became less accurate when adjusting for age, sex, ethnicity, IQ, and country, with the 95% confidence interval including zero effect (B=0.1, 95% CI −0.2 to 0.4, p=0.531). The hallucination dimension was not a robust predictor of draws to decision, whether or not covariates were modelled (unadjusted B= −0.3, 95% CI −0.7 to 0.1, p=0.089). Whereas the CAPE positive symptoms score was negatively associated with the number of beads requested by controls (B= −2.5, 95% CI −3.8 to −1.3, p<0.001), which remained significant even after controlling for age, sex, ethnicity, IQ, and country (B= −1.7, 95% CI −2.8 to −0.5, p=0.006).

## Discussion

The present study was conducted to investigate the relation between the JTC bias and general cognitive ability in first episode psychotic patients. We used the largest to date incidence sample of FEP patients and population-based controls with available data on the JTC bias to address the question: is the link between the bias and the disorder better explained by the mediation of IQ? Moreover, this is the first study to test the association between JTC and genetic susceptibility to schizophrenia.

We found that IQ is accountable for about 80% of the effect of caseness on JTC. The data in fact indicated that case control differences in JTC were not only partially mediated by IQ but were fully mediated by IQ. In other words, the JTC bias in first episode psychosis is not an independent deficit but is part of general cognitive impairment. Previous studies focusing on different explanations between the JTC bias and psychotic disorders suggested that deluded patients tend to request less information because sampling itself might be experienced as more costly than it would be by healthy people (Ermakova et al, 2018; Moutoussis et al, 2011). Although patients at onset seem to adjust their strategy according to the cost of sampling where clearly stated and show more bias to sample less information in the classic task, general cognitive ability still plays an important role in their decision-making behaviour (Ermakova et al, 2018; Moutoussis et al, 2011). Indeed, the inclusion of general cognitive ability in the relation between JTC and clinical status was reported to substantially decrease or nullify a prior significant association (Falcone et al, 2015; Lincoln et al, 2010; Van Dael et al, 2005). Our results are consistent with these studies and provide the first evidence that the genetic variants associated with IQ are negatively correlated with JTC, therefore endorsing the hypothesis that cognition as endophenotype mediates the relation between the bias and the liability for psychotic disorders.

In our study SZ PRS was not significantly associated with JTC. However, as PRS technique is based on contribution of common SNPs variation to genetic risk, its detection is strictly dependent on statistical power (Smaller et al, 2019; Wray et al, 2014). Although GWAS studies on schizophrenia have already identified a substantial number of genetic variants by increasing sample size (Schizophrenia Working Group of the Psychiatric Genomics, 2014), SZ PRS still accounts for 7% of the variance in schizophrenia liability, as we also found in our sample. Whereas, IQ PRS is based on a far larger GWAS (Savage et al, 2018), explaining as twice the variance as SZ PRS in our study. We note that the effect sizes between the schizophrenia PRS and DTD was similar in size to, but, *contrary to expectation, in the same direction as*, the effect size between the IQ PRS and DTD (but note the former association was non-significant). Another possible explanation lies in the fact that JTC seems to be more associated with the continuous distribution of delusions rather than with schizophrenia per se (Broome et al, 2007; Menon et al, 2006; Moritz & Woodward, 2005; Peters & Garety, 2006; Van Dael et al, 2005). In fact, in a large case-control study on FEP patients carried out by Falcone (Falcone et al, 2015), adjusting for IQ and working memory abolished the relation between JTC and clinical status, but the association between JTC and delusion severity remained. Perhaps building a PRS based on the positive symptom dimension or delusions would be more associated with the bias. However, in line with a recent finding that patients with higher delusion severity may have a tendency to *increased* information seeking (Baker et al, 2019), we found delusions were associated with higher number of beads drawn, although with a small effect size and attenuation after adjustment. Whereas, the direction of the effect of PLE on the number of beads requested by controls was expected (Freeman et al, 2008; Reininghaus et al, 2018; Ross et al, 2015), and more interestingly independent from IQ. Perhaps in the absence of general cognitive impairment, the tendency to JTC contributes to psychotic-like experience independent of general intellectual function, whereas in the context of disorder with general cognitive impairment, the JTC bias may not specifically relate to the pathogenesis of psychosis. Future research is warranted to further address this relation.

### Strengths and limitations

Our study has several advantages. These include the large sample size, the study of patients at the onset of illness, and population controls. There are limitations to our study. Although the sample size is several times larger than any individual previous study, it is still modest by genomic standards. We only included a single trial of the task, as previous research has indicated that the measure of draws to decision is very reliable over trials (Ermakova et al, 2018; Ermakova et al, 2014), to facilitate data collection in this large multi-centre study. It can be argued that only performing one trial using the higher cognitive demanding 60:40 version could capture more miscomprehension of the task rather than truly the bias, resulting in overestimating its presence (Balzan et al, 2012a; Balzan et al, 2012b). However, the beads task employed in the study included a practice exercise before the trial. Moreover, the difference between means of beads requested by cases and controls in our study was smaller than reported in a meta-analysis on 55 studies (1.1 vs. 1.4 to 1.7) (Dudley et al, 2016). Nonetheless, future research is warranted to explore more specific cognitive mediation, such as working memory and executive functions (Falcone et al, 2015; Garety et al, 2013; González et al, 2018), between JTC and psychosis using more trials of the Beads Task. The inclusion of more trials in future studies would permit a more detailed interrogation of the task (Ermakova et al, 2018; Ermakova et al, 2014). For example, the use of multiple trials allows modelling of latent variables such as the probing of the distinct roles of cognitive noise and alterations in the perceived “cost” of information sampling, which may have partially distinct genetic bases. We did not measure delusions with a symptom severity rating scale, but rather calculated a measure of a delusions factor using an item-response analysis from OPCRIT identified symptoms, so we caution over-interpretation of the analysis of the relation between DTD and delusions within the patient group.

### Implications

Our results suggested that the tendency to jump to conclusions may be associated with psychosis liability via a general cognitive pathway. In fact, improvements on data-gathering resulting in delayed decision making seem to be driven by cognitive mechanisms as shown in Randomised Control Trials comparing Metacognitive Training (MCT) – which JTC is a core module – with Cognitive Remediation Therapy (CRT) in schizophrenia spectrum disorders (Moritz et al, 2014; Moritz et al, 2013). However, psychosis liability does not necessarily translate to other outcomes of interest in psychosis, such as prognosis; for example, Andreou (Andreou et al, 2014) using the same sample as Moritz (Moritz et al, 2013) found that JTC was the only significant predictor of improvement in vocational status at 6 months follow up in terms of probability of regaining full employment status. Similarly, Dudley (Dudley et al, 2013) found that patients who displayed stable JTC after 2 years showed an increase in symptomatology, whereas stable non-jumpers showed a reduction, as did those who switched to be a non-jumper. Although the aforementioned study did not account for cognition, results are in line with the literature showing the efficacy of MCT on delusion severity and general functioning up to 3-year-follow-up (Favrod et al, 2014; Moritz et al, 2014; Moritz et al, 2013), and even in individuals with recent onset of psychosis (Ochoa et al, 2017). Moreover, a recent study (Rodriguez et al, 2018) which followed up the FEP sample reported in Falcone (Falcone et al, 2015), showed that JTC at baseline predicted poorer outcome in terms of more days of hospitalisation, compulsory admissions, and higher risk of police intervention at 4-year follow-up, even after controlling for IQ.

Thus, although JTC bias might be secondary to a cognitive impairment in the early stages of psychosis, JTC could play an important role in both delusions’ maintenance and clinical outcome. Therefore, targeting this bias along with other cognitive deficits in psychosis therapies and interventions might represent a useful strategy to improve the outcome. However, our data suggest that specific interventions to address this particular cognitive bias in terms of psychosis liability may not provide any advantage over and above cognitive remediation of general cognitive deficits.

## Acknowledgments

This study was funded by the Medical Research Council, the European Community’s Seventh Framework Program grant (agreement HEALTH-F2-2009-241909 [Project EU-GEI]), São Paulo Research Foundation (grant 2012/0417-0), the National Institute for Health Research (NIHR) Biomedical Research Centre (BRC) at South London and Maudsley NHS Foundation Trust and King’s College London, the NIHR BRC at University College London, and the Wellcome Trust (grant 101272/Z/12/Z).

## Conflict of interest

The authors have no conflicts of interest to declare in relation to the work presented in this paper.

